# Precipitation seasonality and soil texture interact to shape dryland recovery from severe disturbance

**DOI:** 10.1101/2023.07.14.549081

**Authors:** Tyson J. Terry, Peter B. Adler

## Abstract

Disturbances drive large changes in plant composition and ecosystem functioning in drylands, but current understanding of how recovery following disturbance depends on the environment is limited due to challenges in analyzing effects of disparate disturbances across abiotic gradients. Here we combine remote sensing and field observations across 5600+ km of natural gas pipeline corridors and adjacent undisturbed vegetation to investigate how recovery from a uniform, severe disturbance varies with factors that influence water availability in drylands. We found that NPP recovery often remains incomplete (42% of sites), except in cold precipitation regimes with low precipitation, and recovery of total shrub cover (median timing of 81 years) was faster on fine-textured soils. Locations with quick recovery of shrub cover were linked with a shift in dominant shrub species and incomplete NPP recovery. Our results indicate that drylands with high productivity or shrub cover are most vulnerable to severe disturbance.

## INTRODUCTION

Ecological disturbances, natural or anthropogenic, cause long-lasting changes to vegetation in dryland systems (Allred et al., 2015; Boyd & Davies, 2012) and have been linked to shifts in stable states (Abella et al., 2021). Understanding recovery trajectories following disturbance is necessary for setting realistic restoration goals while also guiding management and conservation of lands that are susceptible to long-term impacts (Chambers et al., 2017). However, great uncertainty exists concerning recovery rates and how they depend on environmental factors in drylands (Stafford Smith et al., 2009). This uncertainty reflects the short-term nature or limited spatial scale of previous disturbance studies (O’Brien et al., 2022).

The lack of spatially extensive disturbance studies in drylands stems from multiple challenges that make it difficult to compare impacts of different disturbances occurring in separate locations. Disturbances vary in type, severity, timing, and patchiness, all of which influence post-disturbance dynamics and complicate spatial comparisons (Bartels et al., 2016; Diaz et al., 2003). Furthermore, vegetation types characterized by different disturbance regimes usually have different climates. Funding often limits the spatial and temporal scale of recovery monitoring such that disturbance data is often post-hoc, comprised of many space-for-time substitutions that may not capture long-term recovery pathways. Many studies compare the value of a response variable after the disturbance to a pre-disturbance baseline, but this approach may also be unreliable because it assumes a steady-state equilibrium that is unlikely in an era of changing climate and invasive species (Parker & Wiens, 2005; Monroe et al., 2022).

Recovery, which we define as the return of a variable to its undisturbed state following a perturbation (Chambers et al., 2019; Folke et al., 2010; Oliver et al., 2015), comprises two processes: the initial, short-term response of a variable to the disturbance, and subsequent changes over time following the initial response (Hodgson et al., 2015; Ingrisch & Bahn, 2018). These two processes are comparable to resistance (degree of initial response) and engineering resilience (rate of recovery after initial response) (Ingrisch & Bahn, 2018; Nimmo et al., 2015).

A common hypothesis is that the rate of recovery of plant communities following a disturbance depends on resource availability (Chapin et al., 1996; Ridolfi et al., 2006; Shriver et al., 2018; Tilman, 2016). In drylands, water availability is the key resource driving net primary production and vegetation dynamics (Chambers et al., 2014; Davis et al., 2000; Jordan et al., 2020), with previous studies showing that wetter areas or weather periods promote recruitment success (Knutson et al., 2014; Nelson et al., 2014; O’Connor et al., 2020). However, recovery of a system following disturbance requires a return to current undisturbed levels of function and structure (Chambers et al., 2019; Folke et al., 2010; Oliver et al., 2015), and many of the studies supporting this hypothesis in drylands do not compare post-disturbance growth with contemporary undisturbed controls.

An alternative hypothesis is that recovery in drylands may be faster in low resource environments with sparse vegetation. For example, dry areas might recover more quickly than wet areas due to the small amount of growth needed to recover to the relatively low level of productivity found in undisturbed control sites, whereas wetter areas may require substantially more time to recover the comparatively high levels of productivity found in undisturbed sites. This relative-recovery hypothesis emphasizes the importance of undisturbed reference conditions to quantify recovery. Moreover, including undisturbed controls becomes necessary to compare recovery across climatic gradients (Parker & Wiens, 2005), especially in drylands where primary production potential and species composition are strongly tied to water availability (Sala et al., 2012).

Recovery from disturbance often depends on recruitment, which in drylands is tightly linked with soil water availability (Bradford et al., 2019; Maestas et al., 2016; Nelson et al., 2014). Water availability in drylands is largely determined by interactions between precipitation quantity, seasonality, and soil texture (Loik et al., 2004; Maestas et al., 2016; Renne et al., 2019). Coarse soil texture may benefit deep-rooted plants in arid conditions, where high infiltration reduces loss of soil water to evaporation (Maurer et al., 2020b; Noy-Meir, 1973; Sala et al., 1997; Walter, 1964). However, the benefits of coarse soil texture decline with increases in total precipitation, as losses of soil water to leaching outweigh the benefits of low evaporative losses (Noy-Meir 1997). This idea, commonly known as the inverse texture hypothesis, is largely supported by studies of net primary production (Epstein et al., 1997; Sala et al., 1988) and mature plant abundance (Ji et al., 2019; Renne et al., 2019). However, the effect of coarse soils on recruitment remains unclear for shallow-rooted seedlings (Barnard et al., 2019; Boyd & Davies, 2012; Coffin & Lauenroth, 1994). Shifts in precipitation timing from cold to warm seasons can also increase evaporative losses (Lauenroth & Bradford, 2012) and likely increase the benefits of coarse soil texture for plant growth (Renne et al., 2019). Testing hypotheses about water availability and recovery following disturbance in drylands therefore requires careful conservation of interactions between precipitation regimes and soil properties.

Natural gas pipeline corridors provide an opportunity to investigate variation in recovery across environmental gradients, overcoming common limitations of traditional disturbance studies. Pipeline corridors create a near-uniform pulse disturbance that runs hundreds of kilometers, spanning broad soil and climate gradients. Pipeline installation consists of removing all aboveground biomass via bulldozer in a strip we refer to as the corridor (up to 35 m wide), followed by digging a trench and burying a 20-30 cm diameter pipe down the center of the cleared corridor. This disturbance not only removes all plants, but displaces and compacts surface soil (Desserud et al., 2010; Shi et al., 2014). Following construction, topsoils are spread back over the corridor and then seeded for restoration. Post-construction seeding efforts generally use seed mixes to match native species and functional types, but often produce little success and mixed effects on species composition (Farrell & Fehmi, 2018; Rottler et al., 2018). Construction effects are largely concentrated within the pipeline corridors, providing undisturbed neighboring vegetation and soils as a control to measure recovery.

Here we studied two dimensions of ecosystem recovery following pipeline disturbance: total shrub cover and net primary production (NPP). Shrub cover represents a dominant plant functional type in North American drylands (Peinado et al., 1995), and many imperiled wildlife species are considered shrub obligates (Suring et al., 2005). NPP represents the rate at which energy enters the ecosystem and is an indicator of dryland degradation (Wessels et al., 2008; Zika & Erb, 2009). We used annual remotely-sensed estimates of NPP and shrub cover along natural gas pipeline corridors to answer two research questions: (1) How long does it take for NPP and shrub cover in drylands to recover following a disturbance that removes all biomass and disrupts the surface soil? (2) How do mean annual precipitation, precipitation seasonality, and soil texture interact to influence time to recovery of NPP and shrub cover in drylands? We hypothesized that interactions between climate and soils that increase water availability could either a) increase recruitment and speed up recovery of both NPP and shrub cover, or b) create high undisturbed values of NPP and shrub cover that require more time to recover.

## METHODS

### Pipelines

We used The National Pipeline Mapping System (USDOT) to identify pipelines within the Great Basin, Mojave, Chihuahuan, and Sonoran deserts. We selected wide (25+ m) and long (>300 km) pipelines to allow use of high-resolution satellite imagery products derived from Landsat satellites and to span broad, spatial environmental gradients within and among desert systems. Our data came from four natural gas pipeline corridors: Kern River Pipeline (2702 km, built in 1992), Ruby Pipeline (1090 km, built in 2011), El Paso Natural Gas Pipeline (1040 km, built in 1946), and Northwestern Pipeline (860 km, built in 1960). We determined date of construction (initial disturbance) as the midpoint date between the start and completion of construction, a process of 1-2 years. We also contacted pipeline company restoration specialists to confirm there were no additional large-scale disturbances such as herbicide application, new pipelines in the same corridor, or removal of biomass. Other historic disturbances such as grazing and/or wildfire were not accounted for; we assume they have equal effect on disturbed and undisturbed pixels due to their close proximity.

We used annual remotely-sensed metrics from the period 1986-2019 and combined pipelines with differing initial construction dates to quantify recovery up to 73 years since disturbance (Fig. 1). To define the center of pipeline corridors, we manually drew spatial reference lines within Google Earth Engine (Gorelick et al., 2017) using their high-resolution satellite basemap (<1m resolution) while also consulting maps from The National Pipeline Mapping System (USDOT). We selected Landsat pixels (30m resolution) that fit within these pipeline corridors by only using pixels whose centroid was within 3 meters of the reference lines. This allows a maximum of 11% of a pixel to fall outside the pipeline corridor, depending on pixel orientation and position. For undisturbed controls, we used the nearest neighbor along an undisturbed line adjacent to the pipeline corridor. We manually drew the undisturbed comparison lines parallel and near the pipeline corridor (average distance of 120 meters from pipeline corridor). Undisturbed comparison lines were visually inspected with elevational raster datasets and high-resolution imagery to ensure they represented similar topography and land use as the pipeline corridor. Locations where the pipeline corridor or comparison line differed in topography or land use were excluded from the dataset. Both our disturbed and undisturbed reference lines exclude croplands, roads, and urban areas.

**Figure 1.**
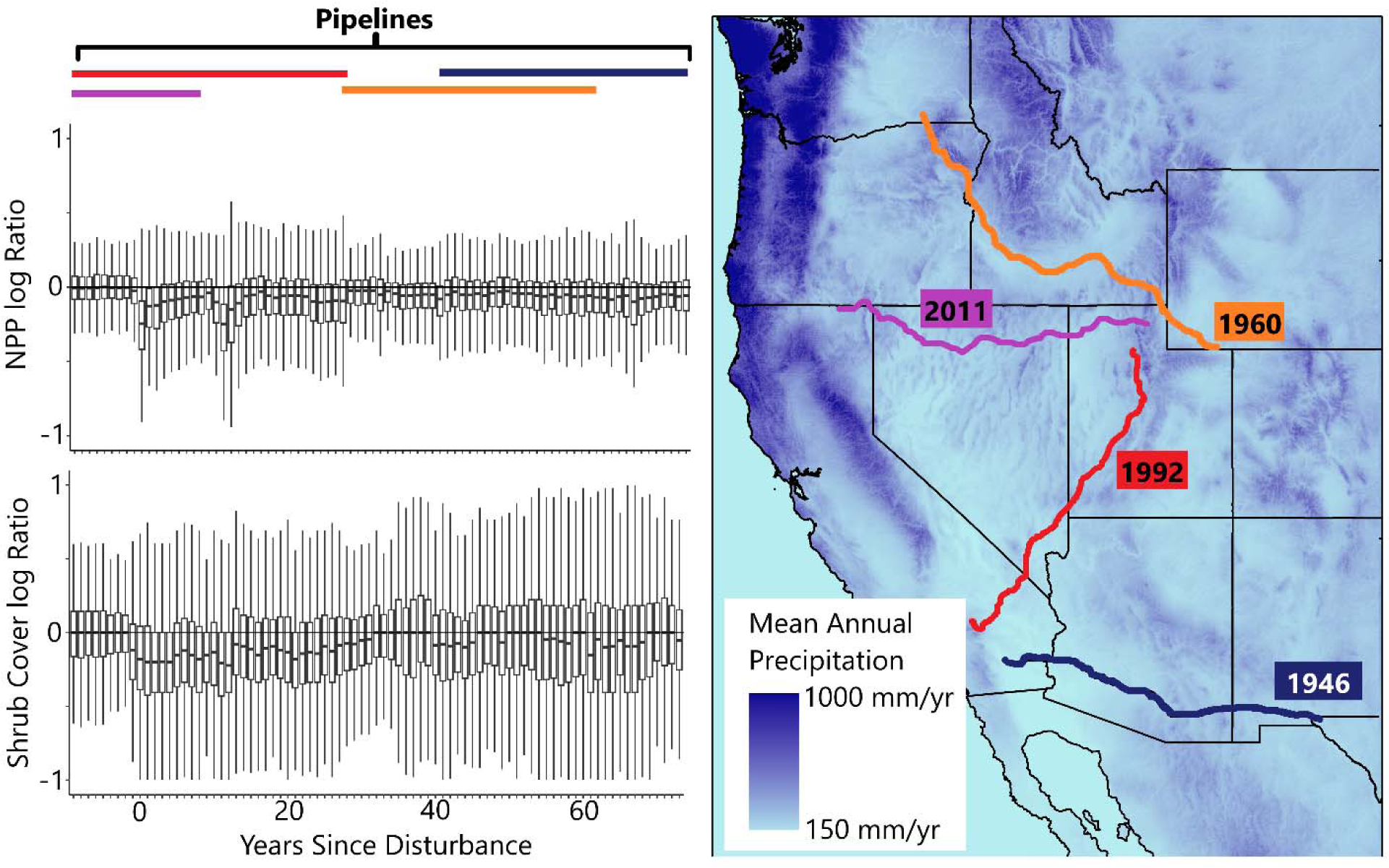
Shrub cover and net primary production ratios before and after pipeline disturbances and pipeline locations. Panel A: shrub cover from remotely sensed data in ratio form of shrub cover recovery (value of 1 indicates equal shrub cover on disturbed pipeline pixel and undisturbed control pixel) with colors indicating the different pipelines used in the study. Panel B: Remotely sensed values of Net Primary Production (NPP) in ratio form (disturbed pipeline corridor value/undisturbed control value). Panel C: Pipeline corridor locations and the year of construction for each respective pipeline. Boxplots in Panel A & B represent the median (center mark), the 25 and 75 percentiles (upper and lower limits of box), and the additional variance of non-outlier data beyond those percentiles (lines extending from boxes).

### Remote Sensing Data

We used remotely sensed estimates of annual NPP data and annual fractional vegetation cover data. Robinson et al. (2018) adapted the MOD17 algorithm, which estimates NPP from MODIS imagery (500m resolution), for use with Landsat imagery (30m). The MOD17 algorithm utilizes photosynthetic active wavelengths and meteorological inputs (short wave radiation, daily minimum and maximum temperature, and vapor pressure deficit) to calculate light use efficiency and scale rates of respiration. Cover type equations were originally estimated from landcover classification to impose biome-specific restraints on respiration and light use efficiency given expected vegetation, but version 2.0 (Jones et al., 2021) uses vegetation type and cover estimated from the rangeland analysis dataset (Allred et al., 2021) to account for within-pixel heterogeneity of plant functional types. These recent developments create NPP estimates that correlate well with 16,591 National Resources Inventory plot-level herbaceous biomass estimates (r=0.63), a comparison made using conversion of the remotely sensed NPP values (Jones et al., 2021). The NPP data represent net carbon uptake (in grams per square meter) on an annual basis.

We used the Rangeland Analysis Platform dataset (version 2.0) for annual estimates of plant cover (Allred et al., 2021). The dataset uses a temporal convolutional neural network to estimate annual percent cover of the following plant functional groups: annual forbs and grasses, perennial forbs and grasses, shrubs, trees, and bare ground. Version 2.0 uses a learned multitask model that examines and learns from interactions of all output variables to improve learning efficiency and prediction accuracy (Caruana, 1998). This dataset utilizes both temporal and spatial smoothing to produce a continuous dataset despite gaps in imagery time-series caused by cloud cover. General prediction error for shrub cover estimates were 6.2 mean square error and 8.8 root mean square error on an out-of-box dataset (Jones et al., 2021).

Mean annual precipitation (MAP) was derived from a gridded climate product, Daymet (Thornton et al., 2022). Daymet produces gridded surfaces (1000 m^2^ spatial resolution) of daily weather parameters based on daily meteorological observations. Annual precipitation values were calculated for each water year (October 1^st^ - September 30^th^) to understand how water inputs relevant to each growing season are impacting recovery. Soil texture data (percent sand content) represents an average value of soil texture estimates of the top 30cm obtained from OpenLandMap (250 m resolution) (Tomislav, 2018). Our metric of precipitation seasonality was mean temperature of wettest quarter (1 km resolution) (Hijmans et al. 2005)

### Data Cleaning

The continuous nature of pipeline corridors and mountainous terrain led to incorporation of many pixels that do not fall within our focus on dryland vegetation. As a result, before initial analysis, we removed all pixels with >20% tree cover and >600 mm of rainfall. Due to the sensitivity of our recovery ratio metric to zeros and extremely small values, we also removed pixels from barren locations with extremely low 30-year mean annual NPP values (<10 g*C/m^2^), and all pixel-years with values of zero. For quality control of remote sensing products, we excluded pixels with >15% temporal gaps caused by missing data and/or cloud cover in their annual Landsat imagery composites.

### Field Surveys

The purpose of these surveys was to provide a qualitative comparison of remote sensing data to field survey data. We also sought to determine if shrubs recovering after the disturbance belonged to the same dominant shrub species as undisturbed controls and if that varied across space, which we could not deduce from our remotely-sensed plant functional group data.

We conducted field surveys at a total of 49 sites across the four pipelines during the years 2021-2022. We specifically selected sites to represent the range of annual precipitation across each pipeline. Visits occurred during peak green-up for each system: mid-April 2022 for the Mojave Desert sites, mid-September 2021 for the Chihuahuan desert sites, and late-May 2021 for the Great Basin sites. At each site we completed four total transects: two parallel 30-meter transects within the disturbed pipeline corridor and two parallel 30-meter transects in undisturbed vegetation 50 meters adjacent to the pipeline corridor. We used a line transect technique to measure percent canopy cover of individual plant taxa. We identified all shrubs to species, with all other taxa to species or genus. For small plants (canopy diameter less than 100 cm), we classified cover as canopy gaps less than 5 cm. For large plants (canopy diameter greater than 100 cm), we classified cover as canopy gaps less than 10 cm.

### Statistical Approach

Our modeling approach assumes that the pulse disturbance of pipeline construction has an initial impact on net primary production and shrub cover, which is followed by more gradual changes over time in the subsequent years since disturbance (YSD) (see equation in Fig. 2). We quantified recovery, our response variable, as the ratio of disturbed to undisturbed values for NPP and shrub cover (remotely sensed) (Fig. 2). With this ratio approach (Avirmed et al., 2015), a value of ∼1, or 0 on the log scale, indicates identical values of a variable in the disturbed and undisturbed pixels. We log-transformed the ratio to normalize the asymmetrical shifts in ratio values that accompany changes in the numerator and denominator values (Isles, 2020). Model predictions were back-transformed to arithmetic scale for ease of interpretation. We used a mixed effects linear model in the lme4 package in R (D. Bates et al., 2015); R Core Team 2022) that assumed the recovery ratio is explained by factors that modify water availability: mean annual precipitation (MAP; Condon et al., 2011), mean temperature of wettest quarter (MTWQ; Nelson et al., 2014), percent soil sand content (Sand; Germino et al., 2018; Maurer et al., 2020a; Renne et al., 2019), and their respective two- and three-way interactions. Each of these environmental covariates varies spatially among pixels (x). We also included a binary disturbance variable, D, with values of 1 (disturbed) or 0 (undisturbed), as well as a variable allowing recovery following disturbance (ln(YSD_x,t_+1)) which varies across pixels (x) and years (t) (Equation in Fig. 2). This time since disturbance variable, (ln(YSD_x,t_+1)), was calculated at all disturbed pixels, with 1 being added to the years since disturbance such that for the year of pipeline completion (YSD = 0), the value would equal 0. As shown in Figure 2, this model allows MAP, MTWQ, Sand, and their interactions to influence the recovery ratio by interacting with the initial impacts of disturbance (D) and years since disturbance (ln(YSD_x,t_+1)).

**Figure 2.**
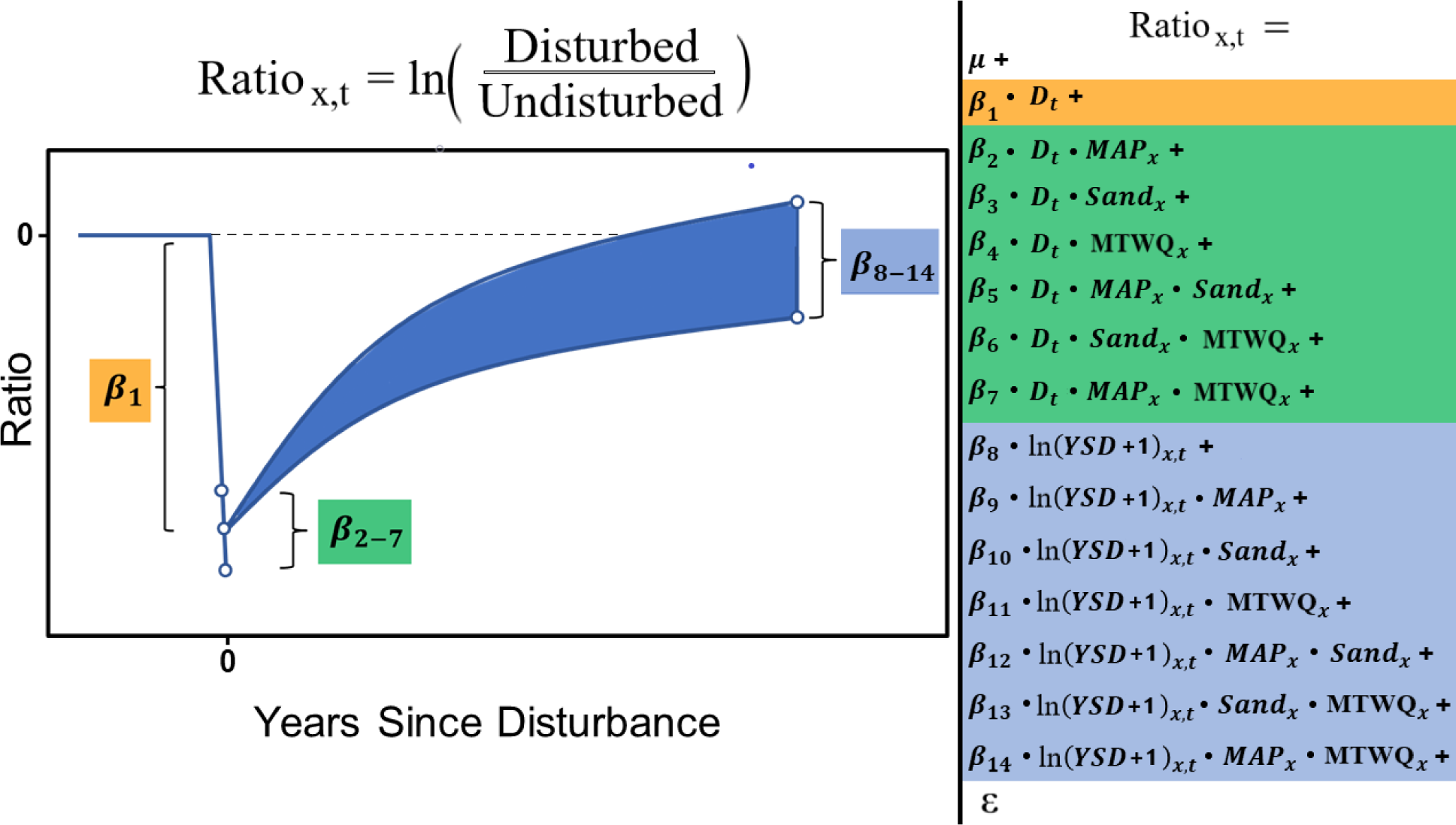
Statistical model visualized to show how covariates can shape initial impact (D) and recovery trajectory (ln(YSD+1)). The yellow color coding indicates the initial impact of disturbance (D). Green color coding indicates how environmental factors of mean annual precipitation (MAP), soil sand content (Sand), Mean Temperature of Wettest Quarter (MTWQ), and their interactions influence the initial impacts of disturbance (D). Blue color coding indicates time since disturbance (ln(YSD+1)) and how environmental factors of MAP, Sand, MTWQ, and their interactions shape long-term recovery. The random effects of pipeline company interacting with both D and ln(YSD+1) are not shown in the formula but were included in our models.

To account for variation in restoration efforts and differing widths of corridors that result from different pipeline companies, we included pipeline identity (e.g. Kern River pipeline) as a random effect on both initial disturbance impact and recovery rate. To account for spatial autocorrelation in our remotely sensed data, we utilized a modified spatial block bootstrap approach (Zhu & Morgan, 2004), where we took random stratified subsamples of our dataset using spatial covariates of MAP and soil sand content as criteria. This approach limits similarity in spatial attributes that occur when pixels occur closely in space, but retains a large range of environmental values to inform the interaction terms within our model. We fit our linear mixed effects model on each stratified bootstrap subsample of the data (∼1300 sites/pixels) that comprised 7% of the dataset and included all years of data for those selected sites (∼37,000 site/pixel-years). We then calculated the median coefficient estimates from 1000 bootstrap iterations and their respective 95% confidence intervals. We considered effects to be statistically significant if the 95% confidence intervals did not overlap with zero. For interpretation purposes and to account for observation error, we quantified full recovery as a return to within 95% of undisturbed levels. Model fit was assessed by checking diagnostic plots to ensure no trends between residuals and fitted values and residuals had a normal gaussian distribution (Figs. S1 & S2).

We created a separate linear regression model to analyze our field data and determine if the shrub species that grows following disturbance matched the dominant shrub species found in undisturbed neighboring vegetation. The response variable was the ratio of dominant shrub species cover, with the cover (%) of the most abundant shrub species in the undisturbed transects as the denominator and the cover (%) of that same species within the disturbed pipeline corridor as the numerator. Explanatory variables were temperature of wettest quarter (described below) and years since disturbance, and we did not include random effects.

### Precipitation Regimes

Our models used the variable “mean temperature of wettest quarter” to represent differences in the vulnerability of precipitation to evaporative demand. To facilitate interpretation and visualization of model results and predictions, we divided the study region into three categorical precipitation regimes based on the temperature of the wettest quarter (Fig. S3). These regions represent areas that receive the majority of their precipitation in cold (<4.5 °C), cool (4.5-17 °C), and warm temperatures (>17 °). Cutoff points for separating these three desert types were determined visually by natural breaks in the distribution of the temperature of the wettest quarter (Fig. S3). More traditional classifications divide deserts into warm deserts (majority of precipitation received during summer) and cold deserts (majority of precipitation received during winter), but the temperature of the wettest quarter provides more information about evaporative demand when most precipitation falls, which was our primary interest.

## RESULTS

### Model Performance

NPP and shrub cover recovery varied dramatically in space and time in our remotely-sensed dataset (Fig. 1). Average R^2^ values after 1000 bootstrap iterations for our NPP and shrub cover models were 0.30 and 0.41 respectively. The substantial unexplained variation in our dataset indicates the importance of sources of variation not included in our model. Both the NPP and shrub cover models indicated significant relationships between our covariates and recovery, with large differences in variable importance between the two models (Table 1).

**Table 1.** Coefficient estimates from bootstrapped linear model explaining recovery of shrub cover and NPP. Lower and upper 95% values represent the 95% confidence interval of the coefficient estimate across 1000 bootstrap samples. Bolded values indicate values with confidence intervals that do not overlap zero, and are interpreted as statistically significant. MAP is mean annual precipitation (mm/water year), Sand is % soil sand content, MTWQ is the mean temperature of the wettest quarter, D is a binary variable indicating whether a given data point is disturbed (1) or undisturbed (0), YSD is years since disturbance.

|  | Shrub Cover |  |  | NPP |  |  |
| --- | --- | --- | --- | --- | --- | --- |
| Coefficient | lower<br>95% | median | upper<br>95% | lower<br>95% | median | upper<br>95% |
| intercept | -0.039 | **-0.036** | -0.025 | 0.026 | **0.032** | 0.047 |
| D (Disturbance) | -0.29 | **-0.253** | -0.136 | -0.146 | **-0.116** | -0.008 |
| D*MAP | 0.101 | **0.114** | 0.154 | -0.026 | -0.01 | 0.045 |
| D*Sand | -0.032 | **-0.026** | -0.004 | -0.06 | **-0.053** | -0.027 |
| D*MTWQ | 0.094 | **0.109** | 0.167 | 0.013 | **0.029** | 0.084 |
| D*MAP*MTWQ | -0.06 | **-0.051** | -0.024 | -0.023 | -0.013 | 0.019 |
| D*MAP*MTWQ | 0.18 | **0.197** | 0.269 | -0.004 | 0.019 | 0.098 |
| D*Sand*MTWQ | -0.069 | **-0.055** | -0.007 | -0.006 | 0.009 | 0.062 |
| ln(YSD+1) | 0.044 | **0.053** | 0.084 | -0.001 | 0.008 | 0.03 |
| ln(YSD+1)*MTWQ | -0.041 | **-0.037** | -0.025 | -0.015 | -0.011 | 0.003 |
| ln(YSD+1)*Sand | -0.003 | -0.001 | 0.007 | -0.003 | -0.001 | 0.007 |
| $\ln(\text{YSD}+1)*\text{MTWQ}$ | -0.03 | **-0.025** | -0.007 | -0.022 | -0.016 | 0.003 |
| $\ln(\text{YSD}+1)*\text{MAP}*\text{Sand}$ | 0.007 | 0.01 | 0.017 | 0.001 | **0.003** | 0.013 |
| $\ln(\text{YSD}+1)*\text{MAP}*\text{MTWQ}$ | -0.053 | **-0.047** | -0.031 | -0.012 | -0.007 | 0.014 |
| $\ln(\text{YSD}+1)*\text{Sand}*\text{MTWQ}$ | 0.002 | **0.007** | 0.021 | -0.002 | 0.003 | 0.016 |

### Initial Impacts on NPP

The initial impact of pipeline disturbance decreased annual NPP 11% on average. Sites with coarse soil were more negatively impacted, whereas areas with warmer precipitation regimes were less negatively impacted (Table 1, Fig. 3). Interactive effects of MAP and precipitation seasonality on initial impacts were large but variable (Table 1).

**Figure 3.**
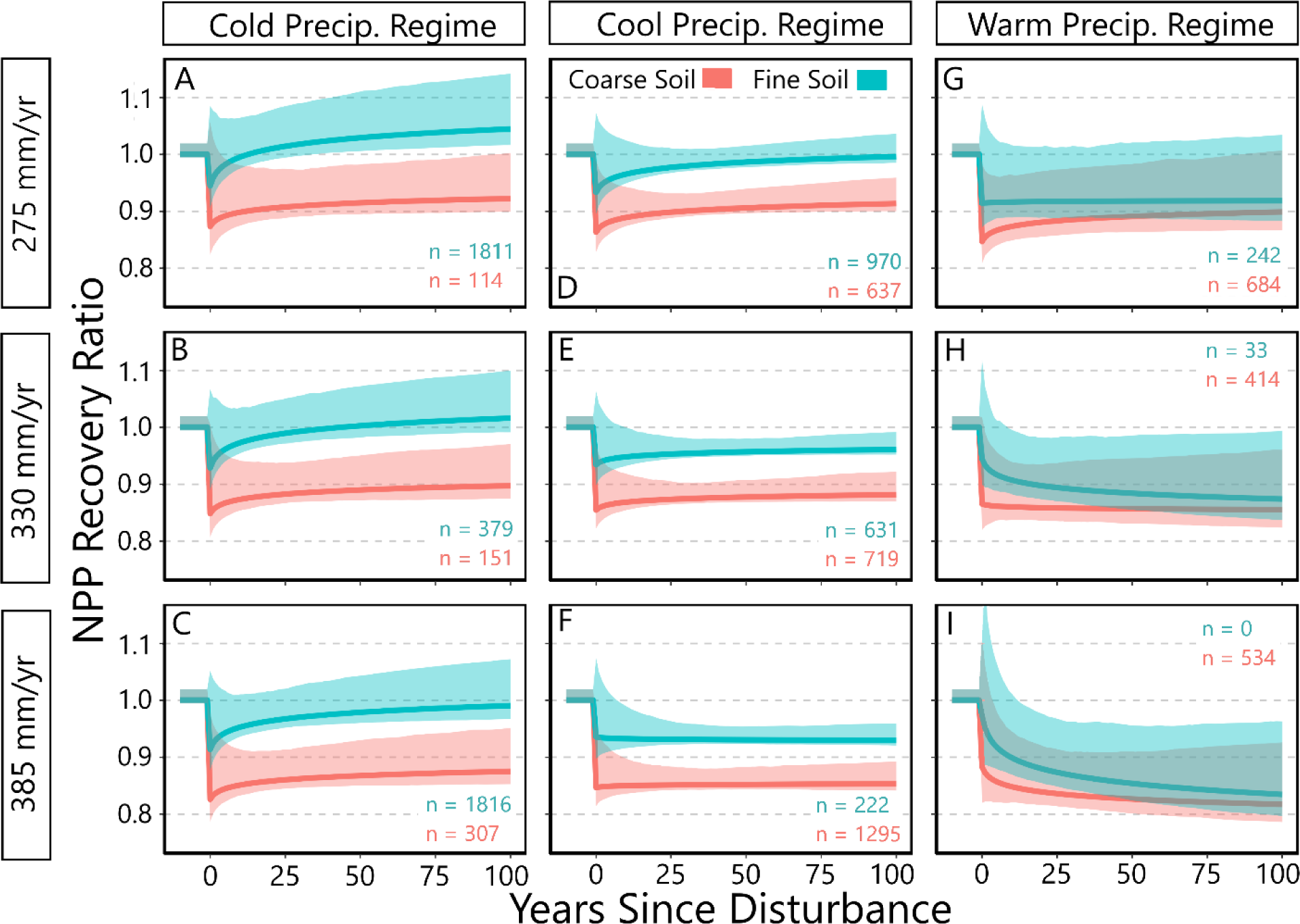
Predicted patterns of net primary production (NPP) recovery across North American deserts. The three columns represent spatial differences in precipitation timing: areas that receive a majority of their precipitation during periods of cold, cool, and warm temperatures. The three rows represent different scenarios of mean annual precipitation (MAP). Colors represent coarse (40% soil sand content) and fine soil texture (30% soil sand content). Recovery values (y-axis) are a ratio of disturbed NPP (numerator) in relation to undisturbed NPP (denominator). A value of 1 indicates equal shrub cover on disturbed sites and undisturbed controls. N count values represent the number of corresponding sites within the dataset used for training.

### Long-Term Recovery of NPP

The long-term value of NPP was largely determined by the initial drop in NPP at the time of disturbance because little recovery occurred except in cold, dry precipitation regimes (∼275 MAP) with fine soils (Fig. 3 Panel A). Recovery rate following initial disturbance impacts generally decreased with warmer precipitation regimes and higher annual precipitation (Fig. 3). The interaction of Sand and MAP was the only interaction in our model that significantly affected recovery rate following initial disturbance (Table 1).

### Initial Impacts on Shrub Cover

Total annual shrub cover was significantly impacted by the pipeline disturbance, dropping 23% on average. MAP, precipitation season, and their interaction had the strongest effects on initial impacts of disturbance (Table 1). Areas that were wetter (higher MAP) and belonged to warmer precipitation regimes had smaller initial negative impacts (Fig. 4 Panels G-I). Effects of coarse soil texture (higher sand content) switched from slightly positive to negative when transitioning from cold to warm precipitation regimes (Fig. 4, Table 1).

**Figure 4.**
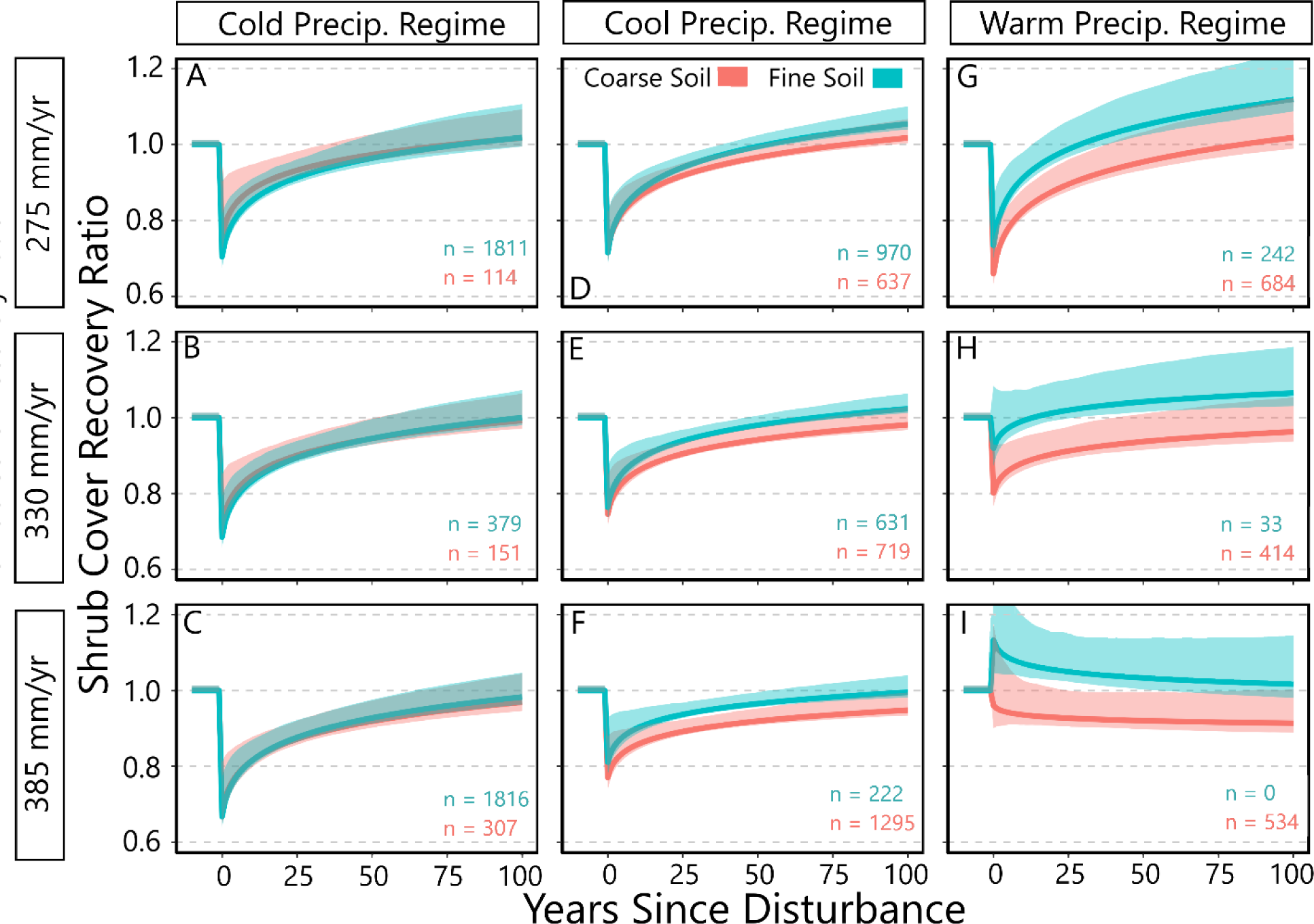
Predicted patterns of shrub cover recovery across North American deserts. The three columns represent spatial differences in precipitation timing: areas that receive a majority of their precipitation during periods of cold, cool, and warm temperatures. The three rows represent different scenarios of mean annual precipitation (MAP). Colors represent coarse (40% soil sand content) and fine soil texture (30% soil sand content). Recovery values (y-axis) are a ratio of disturbed shrub cover (numerator) in relation to undisturbed shrub cover (denominator). With this system a value of 1 indicates equal shrub cover on disturbed sites and undisturbed controls. Margins represent 95 percent confidence intervals from 1000 bootstrap model iterations. N count values represent the number of corresponding sites within the dataset used for training.

### Long-Term Recovery of Shrub Cover

Higher MAP and warmer precipitation season both significantly decreased the long-term shrub recovery rate (Table 1, Fig. 4). Negative impacts of coarse soil on long-term recovery were stronger in warmer precipitation regimes (Fig. 4 Panels G-I). A combination of high MAP and warm precipitation regime (not present in our data) shifted the recovery trajectory from initial-drop followed by positive recovery, into a trajectory of initial increase, followed by subsequent decline (Fig. 4 Panel I).

### General Recovery Timelines

Across all our pipeline sites, our models predicted that 50% of sites will recover shrub cover within 81 years of disturbance, with 79% of sites recovering within 150 years (Figs. S4 & S5), assuming recovery rates follow current trajectories. NPP recovery was predicted to occur rapidly (<20 years since disturbance) at 40% of sites, with almost all other sites unlikely to recover within 150 years after the disturbance (Figs. S4 & S5).

### Field Observations

Field data showed that 52% of sites (n=23) at two old pipelines (61 & 75 years since disturbance) had recovered total shrub cover to current undisturbed levels (Fig. S6), agreeing well with our remotely sensed estimates of 53% recovery of all pipeline pixels for the same pipelines at that time post-disturbance. Average recovery ratio values based on field data across the same pipelines was 1.02, whereas the average of model estimates for the same pipelines was lower at 0.95.

Precipitation regime and years since disturbance explained 38% of variance in dominant shrub recovery ratios across 49 field sites. Our field surveys indicate that shrub species composition following disturbance often differed from the dominant undisturbed shrub species and was significantly affected by MTWQ (df = 44, t score = -3.955, *p* < 0.001). Specifically, the similarity in shrub species composition between disturbed and undisturbed sites was highest in colder precipitation regimes (Fig. S6).

## DISCUSSION

Current understanding of recovery following disturbance has been limited by the spatiotemporal scope of previous studies and complications such as variation in the type, timing and intensity of disturbance events across different plant communities. Our comparison of disturbed and adjacent undisturbed vegetation across spatially extensive pipeline disturbances allowed us to investigate how recovery varied across environmental gradients that influence water availability. Though not representative of any natural disturbance, data from pipeline corridors enabled us to examine recovery across space following a uniform disturbance that removed vegetation and disrupted the surface soil.

We found that the initial impacts of disturbance and the rate of recovery following disturbance vary greatly across time and space (Figs. 3 & 4). NPP and shrub cover did not recover simultaneously, but rather showed opposite patterns in recovery. Disturbance impacts to NPP were not large, but only 42% of sites were projected to recover to undisturbed production NPP levels within 100 years of disturbance (Figs. S4 & S5). In contrast, disturbance impacts on shrub cover were large, but most pixels (61%) were projected to recover within 100 years (Figs. S4 & S5).

Our first research question was, how long does it take for NPP and shrub cover in drylands to recover following a disturbance that removes all biomass and disrupts surface soil? We found that median recovery time for shrub cover was 81 years after the disturbance, with most sites (80%) projected to recover within 200 years after the initial disturbance. NPP recovery was projected to occur at 40% of sites within 20 years, but the majority of the remaining sites were not projected to recover within 200 years (Fig. S4).

Our second research question asked, how do mean annual precipitation, precipitation timing, and soil texture interact to influence recovery of NPP and shrub cover in drylands? Our results indicate that more precipitation does not drive rapid recovery. Rather, the initial impacts of disturbance and rate of recovery depend on soil texture, MAP, precipitation timing, and their interactions. In general, our results indicate that NPP recovery was fastest in cold and dry locations, and long-term recovery potential decreased in locations with coarse soils and warmer precipitation regimes (Fig. 3). Shrub cover recovery was quickest on fine soils in warm precipitation regimes (Fig. 4), but our field studies indicated that this recovery reflects the establishment and growth of different shrub species than those present in neighboring undisturbed plots (Fig. S6).

Both shrub cover and NPP values did not decline to zero following disturbance. This is likely due to several factors. First, we used annual NPP values, so depending on when the disturbance occurred, there is likely production prior to or in the period after the pipeline disturbance, but still within the year of the disturbance. Second, Landsat pixels do not always fit perfectly within pipeline allowing very small portions of undisturbed vegetation within each disturbed pixel. Third, though the remotely sensed estimates of NPP and shrub cover operate on reflectance, the algorithms were trained on large spatial datasets that may limit sensitivity to abrupt changes, and instead focus on predicting average landscape values to minimize error.

### Contrasting Shrub and NPP Recovery

Areas that recovered shrub cover quickly did not recover NPP quickly, and vice versa (Figs. 3 & 4). NPP recovery was concentrated in areas of cold precipitation regimes or very low NPP (Fig. 3). We hypothesize that cold precipitation regimes favor NPP recovery due to high growth of forbs and grasses following disturbance. Our field surveys support these findings, with disturbed pipeline corridors in a cold precipitation regime having on average 2.5-fold more forbs and grasses than neighboring undisturbed vegetation (10 years after disturbance). Whereas we observed little to no grasses or forbs in the disturbed pipeline corridors in the warmer Mojave Desert (30 years after disturbance), a system that sees periodic bursts of annual grasses and forbs during above-average water years. High grass cover has been shown to limit shrub recruitment (Davidson et al., 2019; Germino et al., 2018), particularly in cold precipitation regimes (Bates & Davies 2022), which may explain the quick recovery of NPP and slower shrub cover recovery. However, in warm precipitation regimes, disturbance linked with soil disruption and erosion has been hypothesized to promote a shrub stable state (Okin et al., 2009; Schlesinger et al., 1990).

Our model predicts incomplete NPP recovery across many sites (58%), including those that had full shrub cover recovery (Fig. S5). We hypothesize that this mismatch is tied to changes in species composition. In cold precipitation regimes, our field observations indicate that the same shrub species occur on disturbed and undisturbed sites (*Artemisia tridentata, Sarcobatus spp., Atriplex spp*.), whereas in warmer precipitation regimes recovery of the dominant undisturbed shrub (usually *Larrea tridentata or Prosopis spp.*) was limited (Fig. S6). In place of dominant shrubs such as *Larrea spp.*, disturbed plant communities in warmer precipitation regimes consisted of small early successional species such as *Gutierrezia sarothrae, Ambrosia dumosa,* and small grass species. Many of these early successional species are known to reduce aboveground forage and production of plant communities due to competitive effects on other species, especially during years with below average precipitation (Nagy et al., 2021; Tschirley & Clark, 1961). This supports the findings of a review on disturbance in drylands showing that despite recovery of herbaceous cover, plant species composition often shifts to a new stable state following disturbance (Abella et al., 2021).

### Relative-Recovery Hypothesis

We hypothesized that time to recovery would either increase with higher water availability, or that high water availability would be associated with high undisturbed levels of production or shrub cover that would recover more slowly. We found that despite more production and shrub cover following disturbance, areas with high undisturbed levels of shrub cover and/or NPP did not return to undisturbed levels more quickly than areas with low undisturbed levels (Fig. 5). We observed that some areas with very low undisturbed NPP appear to slightly increase in production following disturbance (Fig. 5). This pattern supports a relative-recovery framework, where time to full recovery may be slower in resource rich locations despite faster year-to-year increases in shrub cover or NPP following disturbance.

**Figure 5.**
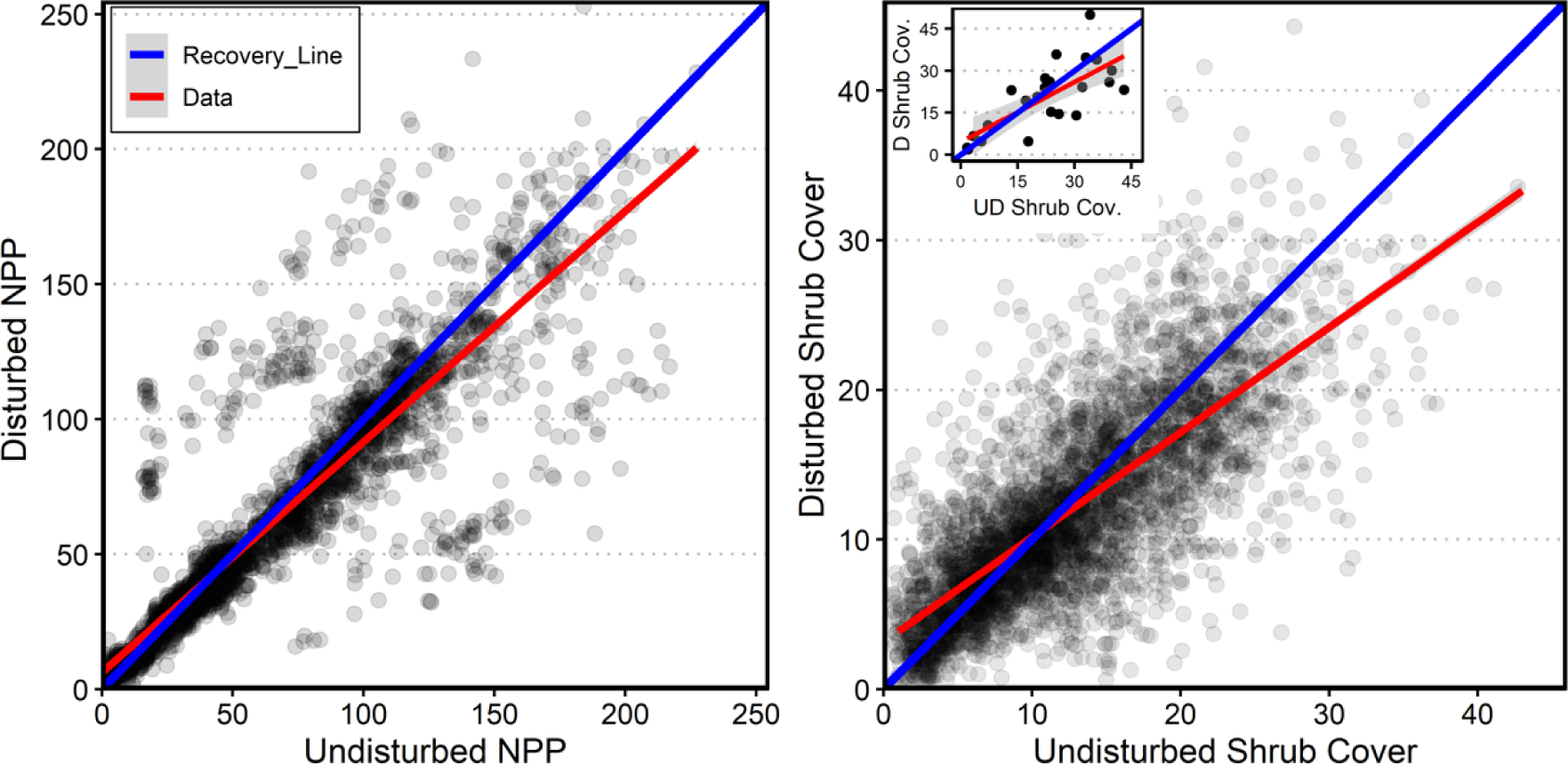
Plots comparing levels of Net Primary Production (left) and total shrub cover (right) at disturbed locations with respect to their respective undisturbed control levels (x axis) 50-55 years after disturbance. Gray dots represent individual sites. The blue line represents recovery, where levels of a variable in a disturbed location return to the level of their undisturbed control. The red line represents a regression line for the dots/sites. Both plots use remotely sensed estimates of pixels disturbed 50-55 years after pipeline construction. The inset graph in the right panel uses field observations of total shrub cover at sites disturbed 61 & 75 years ago. All red regression lines have a gray margin representing standard error, though some margins are similar to the width of the line.

Land use history is an important factor to consider in the relative-recovery framework. Arid locations with low undisturbed values of NPP and shrub cover may be more likely to be degraded than more productive systems due to historic livestock grazing (Hoover et al., 2020). These systems may recover quickly, but to a degraded or early successional state rather than to a later successional state. This emphasizes the importance of understanding pre-disturbance conditions, especially if they vary systematically across space with climate and land-use. Our analysis used undisturbed vegetation as a reference, but assessing the condition of the reference pixels was beyond the scope of our study.

Drylands are complex systems that are sensitive to physical disturbance (Svejcar & Kildisheva, 2017). Areas with high production tend to have more plant species and more diverse soil microbes (Adler & Levine, 2007; Maestre et al., 2015). Our results indicate that the complex biotic interactions that support high NPP and shrub cover within water-limited systems may require more time to reestablish following disturbance. Differential response of low and high productivity areas to disturbance may also be linked to changes in abundance of invasive annual grasses often associated with disturbance (D’Antonio & Vitousek, 1992), which can potentially lead to a higher ephemeral production (Wolkovich et al., 2010). We recommend prioritizing the conservation and restoration of productive dryland plant communities that are unlikely to return to prior levels of function and composition, analogous to biodiversity hotspots in conservation biology.

## Acknowledgments

We would like to thank Sasha Reed, Bill Smith, Scott Ferrenberg, Osvaldo Sala, Brooke Osborne, Steven Lee, and Sam Jordan for their feedback and ideas that strengthened this project. This research was supported by a grant from the US Department of Defense, Strategic Environmental Research and Development Program (grant #201940).

