## Supplemental Material for "Precipitation seasonality and soil texture interact to shape dryland recovery from severe disturbance"

Supplemental Figures

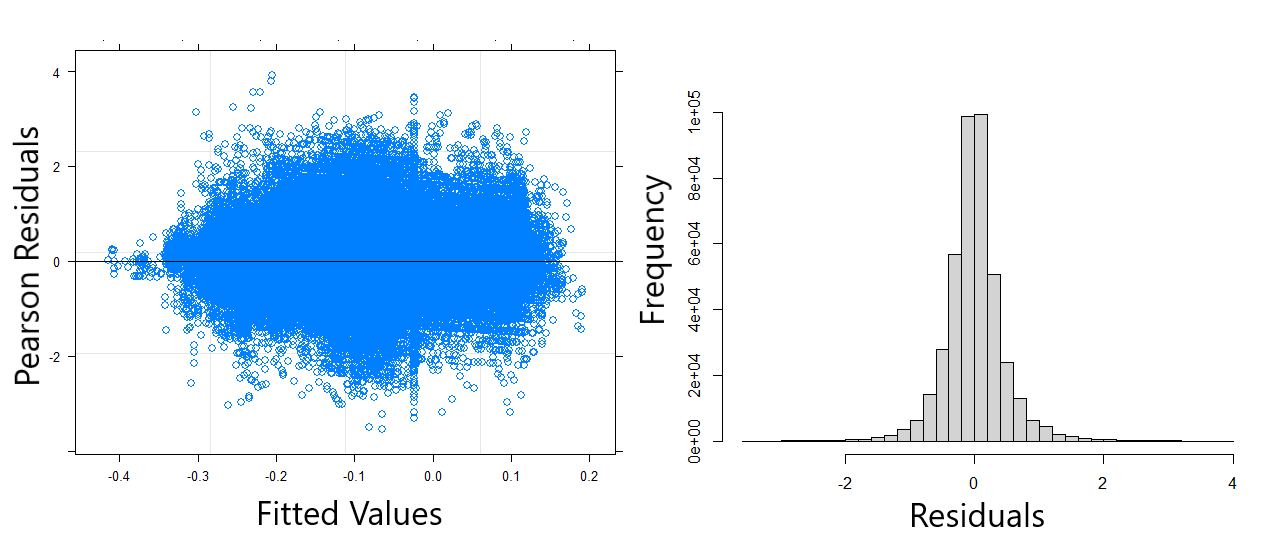

Figure S1. Diagnostic plots of shrub cover model. Residuals versus fitted values (left) and histogram (right).

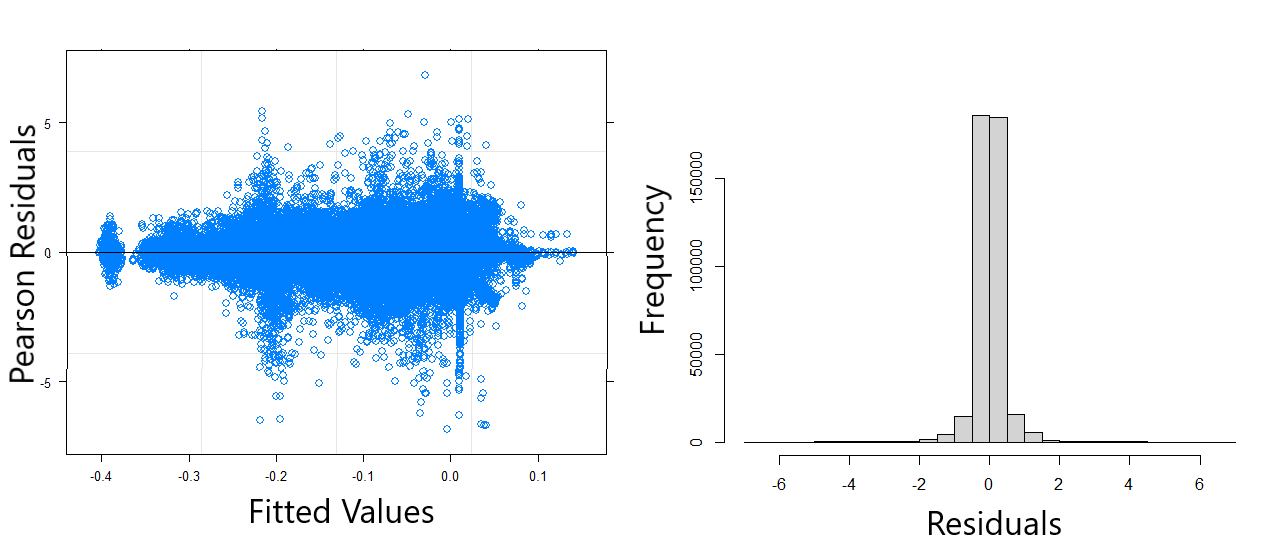

Figure S2. Diagnostic plots of NPP model. Residuals versus fitted values (left) and histogram (right).

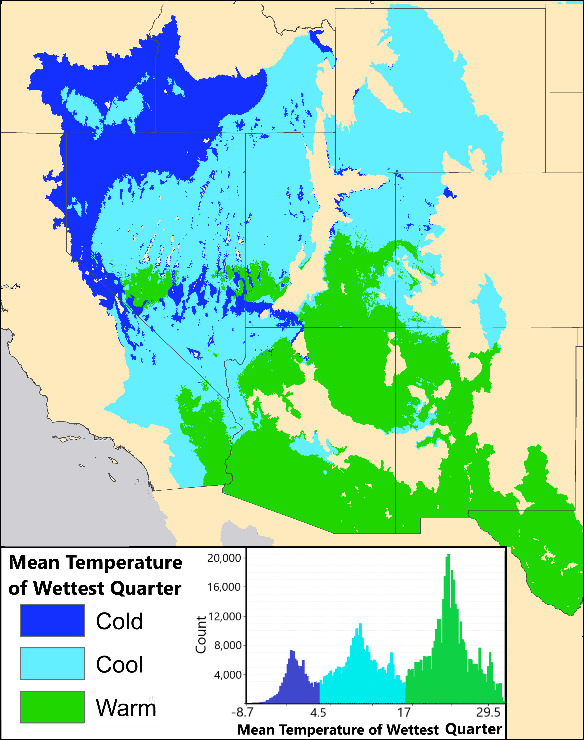

Figure S3. Map indicating the spatial extent of three precipitation regimes in North American Deserts: areas that receive the majority of their precipitation in cold temperatures, areas that receive the majority of their precipitation in the cool temperatures, and those that receive the majority of their precipitation in warm temperatures. Histogram included to show distribution of the temperature of wettest quarter (°C) used to separate the three precipitation regimes.

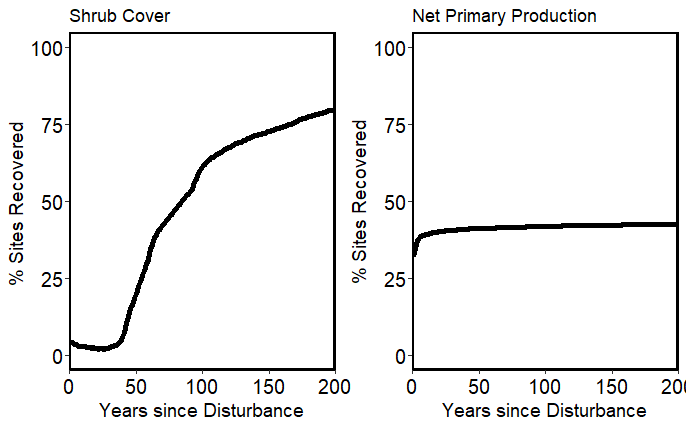

Figure S4. Modeled recovery dynamics of all pixels within study. Recovery is defined here as disturbed areas reaching 95% NPP or shrub cover of concurrent undisturbed controls

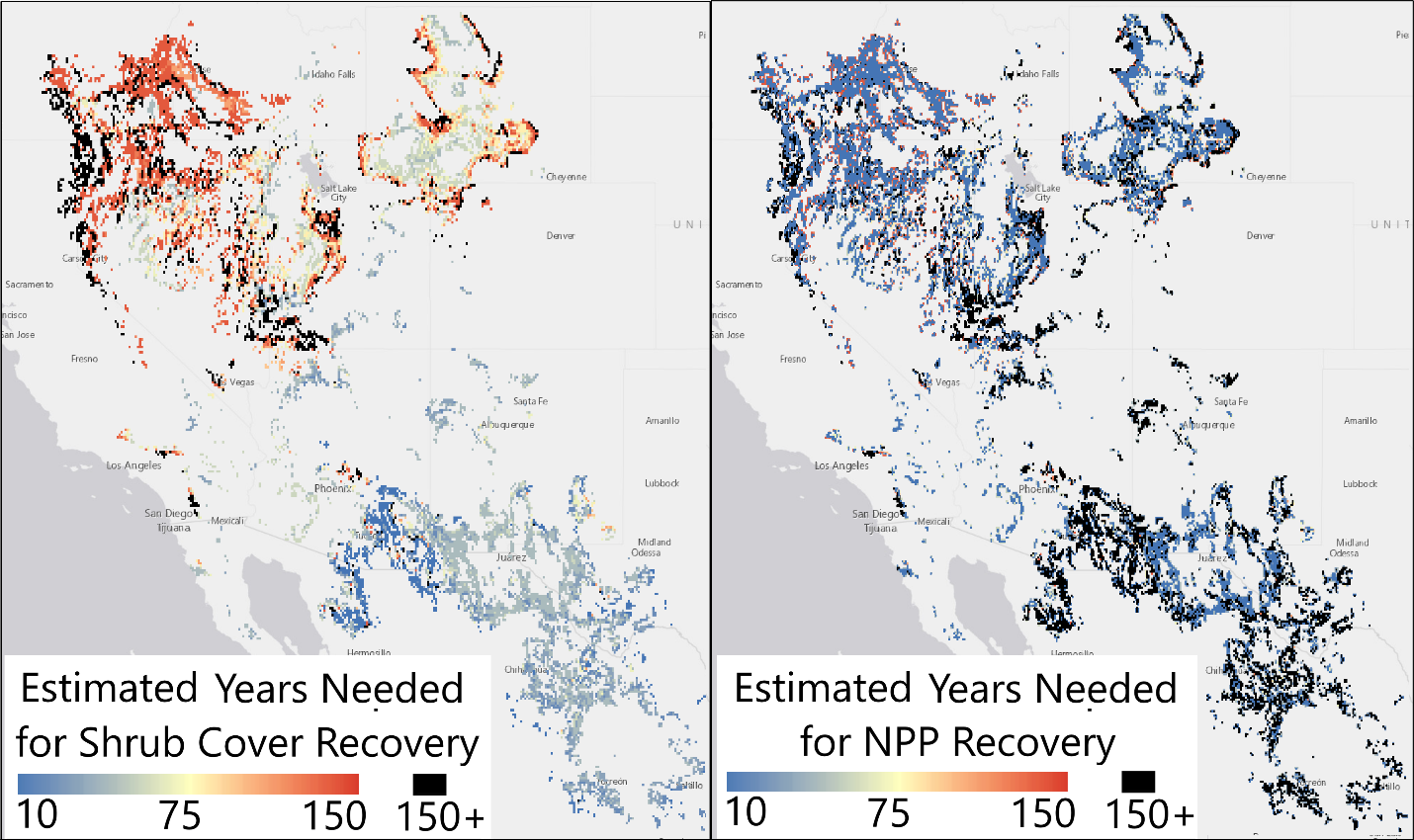

Figure S5. Estimated recovery timing of shrub cover (left) and NPP (right). Only pixels within North American deserts that have similar abiotic conditions (same combination of % sand, mean annual precipitation, and precipitation seasonality) as pixels in our pipeline dataset are mapped.

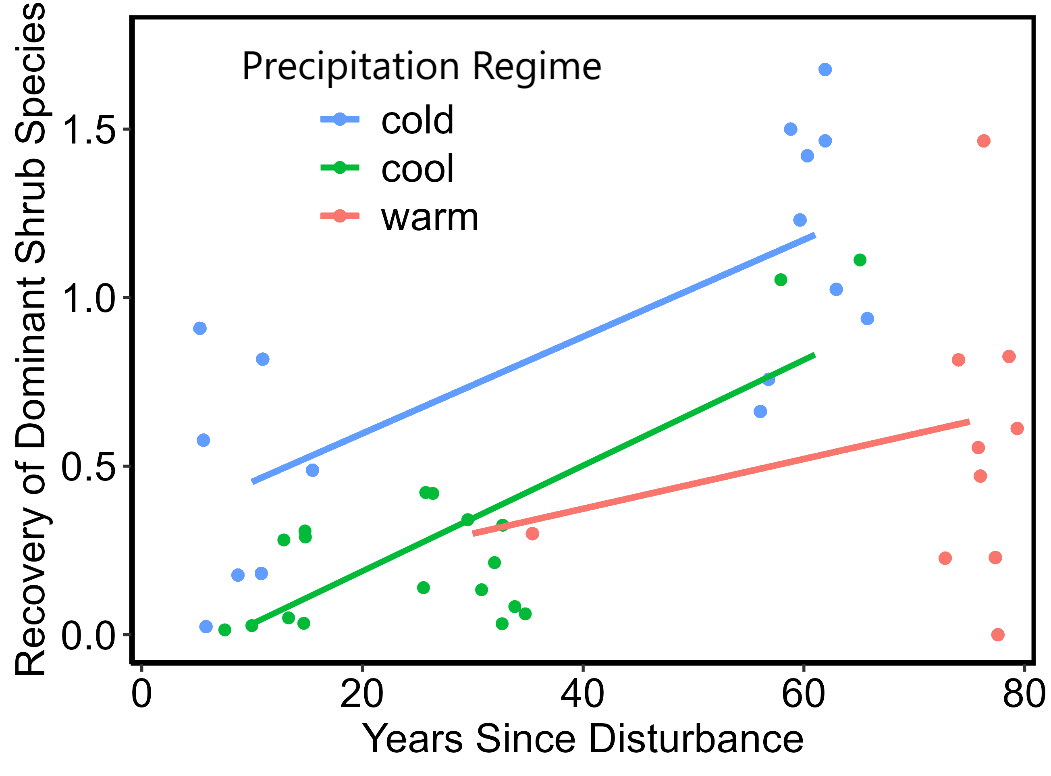

Figure S6. Recovery ratio values (shrub cover in disturbed plots/shrub cover in undisturbed control plots) of the dominant shrub species in undisturbed conditions across our field observations. Colors represent different precipitation regimes: those that receive the majority of their precipitation at cold, cool, and warm temperatures.
